# Female Chemical Warfare Drives Fitness Effects of Group Sex Ratio

**DOI:** 10.1101/100354

**Authors:** Imroze Khan, Arun Prakash, Swastika Issar, Mihir Umarani, Rohit Sasidharan, N. Jagadeesh, Prakash Lama, Radhika Venkatesan, Deepa Agashe

## Abstract

In animals, skewed sex ratios can affect individual fitness either via sexual interactions (e.g. intersexual conflict or intrasexual mate competition) or non-sexual interactions (e.g. sex-specific resource competition). Because most analyses of sex ratio focus on sexual interactions, the relative importance of these mechanisms remains unclear. We addressed this problem using the flour beetle *Tribolium castaneum*, where male-biased sex ratios increase female fitness relative to unbiased or female-biased groups. Although flour beetles show both sexual and non-sexual (resource) competition, we found that sexual interactions did not explain female fitness. Instead, female fecundity was dramatically reduced even after a brief exposure to flour conditioned by other females. Earlier studies suggested that quinones (secreted toxins) might mediate density-dependent population growth in flour beetles. We identified ethyl- and methyl-benzoquinone (EBQ and MBQ) as the primary components of adult stink glands that regulate female fecundity. In female-biased groups (i.e. at high female density), females upregulated quinones and suppressed each other’s reproduction. In male-biased groups, low female density lead to low quinone levels, allowing higher fecundity. Thus, quinones serve both as indicators and mediators of female competition, resulting in the observed fitness decline in female-biased groups. Our results underscore the importance of non-sexual interference competition that may often underlie the fitness consequences of skewed sex ratios.

## INTRODUCTION

Theoretical models predict that a balanced sex ratio is optimal for most diploid species [e.g. 1–4]. However, the adult sex ratio varies substantially both across species and populations, and a large body of research has analyzed the evolutionary consequences of skewed sex ratios. In particular, the impact of sex ratios on reproductive behavior and associated intra- or inter-sexual conflict has received a lot of attention, since reproductive traits are expected to face strong natural as well as sexual selection. For instance, male-biased sex ratios can be detrimental because of direct or indirect male harassment, decreasing female longevity [e.g. 5], fecundity [e.g. 6], or offspring survival [e.g. 7]. More generally, increased male-male competition can result in elevated sexual conflict, reducing female fitness [8]. On the other hand, under a male-biased sex ratio, females may benefit from the opportunity to selectively mate with higher-quality males [9], or benefit from polyandry by increasing offspring heterogeneity [10].

These hypotheses explaining the relationship between adult sex ratio and female fitness largely depend on sexual interactions involving mating and associated behaviors. However, individual fitness is also determined by non-sexual interactions that alter survival and life history [11]. Such non-sexual interactions may be especially important for females, who often compete more strongly for resources (such as food or space) rather than for mates [12]. A growing body of work on mammals suggests that female competitive interactions may be important for many species [13]. For instance, in cooperatively breeding meerkats, dominant females increase their reproductive success by suppressing subordinate females’ reproduction, reducing competition for resources [14]. Female parasitoid wasps face intense intraspecific competition for suitable hosts, and use physiological and chemical mechanisms to determine the outcome in their favour [15]. In such cases, resource rather than sexual competition may govern female fitness. More generally, changes in sex ratio may be confounded with changes in the density of one or both sexes. Hence it is important to quantify the contribution of sexual interactions vs. non-sexual competition on individual fitness.

We tested the impact of adult sex ratio on female fitness in the red flour beetle *Tribolium castaneum*, a global pest of stored grains. Typically, populations of *T. castaneum* exhibit a balanced sex ratio [16]. Both sexes mate multiply and exhibit mate choice [16–20], but the degree and fitness consequences of polyandry vary across populations [21]. Last-male sperm precedence is well known, although there is large variation in male offense capability [22], which may potentially trade off with sperm defense ability [23]. Hence, sexual selection, sexual conflict and intra-sexual competition are important aspects of fitness in flour beetles. In addition, individuals compete strongly for resources, which determines density-dependent population growth rate [24]. Together, these features make *T. castaneum* an attractive model to analyze the impact of different forms of individual interactions on fitness in the context of imbalanced sex ratios.

We tested the consequences of skewed adult sex ratio on multiple proxies of female fitness: female reproductive fitness (egg production and total surviving offspring), lifespan, and immune function. We found that for in each case, females from male-biased groups had higher fitness compared to females from either unbiased or female-biased groups. Neither male intra-sexual competition nor opportunity for female or male mate choice explained this fitness difference. Instead, we found that negative density-dependent interactions between females were responsible for the fitness effects of group sex ratio. We identified benzoquinones as key chemicals mediating these interactions, suggesting female interference competition. Thus, we present a case where fitness consequences of group sex ratio arise from female-driven non-sexual resource competition, rather than sexual interactions.

## METHODS

### Generating experimental individuals and sex ratio groups

We used an outbred laboratory population of *T. castaneum* [25] maintained on whole wheat flour (henceforth “flour”) at 34°C (±1°C) on a 45-day discrete generation cycle. For all experiments, we allowed ~2000 individuals to oviposit in 750 g of wheat flour for 48 h, and collected their offspring at the pupal stage. We housed pupae of each sex separately in 1.5 ml micro-centrifuge tubes (3 pupae of the same sex in 1g flour) for 10 days post-pupation. The pupal stage lasts for 3-4 days; we thus obtained ~7 day old (post-eclosion), sexually mature virgins. We grouped these adults into three sex ratio treatments (*n* = 8-9 groups per sex ratio): (1) Male-biased groups (henceforth “MB”: 3 males + 1 female) (2) Unbiased groups (henceforth “UB”: 3 males + 3 females) and (3) Female-biased groups (henceforth “FB”: 1 male + 3 females). Each group was maintained in a plastic Petri plate (60 mm diameter, Tarsons) at a density of 1 beetle/g flour (chosen to minimize resource limitation), and placed in a 34°C (±1 °C) incubator. Thus, MB and FB groups were given 4g of flour and UB groups received 6g of flour. We chose the UB treatment such that we could determine the effect of skewed sex ratio given the same number of males (UB vs. MB, 3 males each) or females (UB vs. FB, 3 females each); note that multiple males or females were necessary to evaluate mate choice hypotheses. To test whether group size altered the effects of group sex ratio, we set up additional groups with a total of 12 beetles/group (n = 10 groups per sex ratio) but the same sex ratio (FB: 9 females + 3 males; MB: 3 females + 9 males) and density per g flour.

### Measuring fitness effects of group sex ratio

Females in each experimental group could oviposit continuously during the experiment, but adult cannibalism would confound estimates of female fecundity and fertility. Hence, after constituting the groups described above, we estimated the total offspring produced per female by periodically isolating females: every 5 days for the first 30 days, every 7 days for the next 28 days, and then every 14 days for another 42 days. Thus, we obtained estimates of female reproductive fitness as a function of sex ratio as well as age, until females were 102 days old. During each fitness assay, females oviposited individually for 24 hours in plastic Petri plates (60mm diameter, Tarsons) containing 5g flour, after which they were placed again into their original group and provided fresh flour. Thus, offspring from each female developed independently in the oviposition plates. After three weeks, we counted the total number of offspring in each plate (including larvae, pupae and adults) as a proxy for the total reproductive fitness of each female. To separate female fecundity (egg production) from fertility (egg survival), we counted the number of eggs laid by 21-day-old females after the 24-hour oviposition period as a direct measure of female fecundity. We also monitored mortality of individuals from each group, and tested proxies of immune function (see SI methods for details).

### Quantifying mating behaviors

To quantify mating-related behaviors as a function of group sex ratio, we set up additional experimental groups using virgin adults (*n* = 15-16 groups per sex ratio). To identify females in FB and UB groups, we used a paintbrush to mark the elytra with a small dot of non-toxic acrylic (model master) paint, using one of three colours (yellow, green or red). To control for any effects of the paint on mating behavior, we marked 1/3^rd^ of the MB females with each colour. Thus, all females in the experiment were marked regardless of the sex ratio of their group. After marking the elytra, we placed females individually in wells of a 96 well microplate and allowed the paint to dry for 6 hours. Since we wanted to quantify mating behaviors directed towards females, we did not mark males. We distributed marked females into sex ratio groups as described in the previous section. A week later, we separated individuals from the flour, placing each group in a 60 mm Petridish whose bottom was covered with filter paper to allow beetles traction for walking. We observed each group for 30 minutes, noting the number and timing of matings received per female. A mating event was recorded when the male mounted the dorsum of the female and attempted or achieved copulation. We quantified (1) total number of copulatory mounts (matings) (2) total time spent in copula (3) average copulation time and (4) mating latency (time until the first copulation). Immediately after the behavioral assay, we separated females and measured their reproductive output (as described above) to test whether female fitness was correlated with mating behaviour.

### Testing the effect of conditioned flour on female fitness

Flour used by *Tribolium* beetles gets “conditioned” by the accumulation of secreted quinones, pheromones, and excreta. We tested whether flour conditioned by FB and MB groups differentially affects the fitness of females exposed to the flour. We allowed FB and MB groups (constituted as described earlier) to condition 4g of flour for 10 days. After this, we removed the adults, froze the flour at −80 °C for 15 minutes and sifted it through a fine mesh (300 μm pore size, Daigger) to kill and remove juvenile stages. We placed a virgin male and a virgin female (both 7 day old) in 2g of this conditioned flour for 6 hours, 1 day or 4 days (*n* = 12 pairs per conditioning treatment per duration of exposure). Following this, we isolated the female from each pair and allowed her to oviposit for 24 hours in fresh flour. The next day, we counted the number of eggs laid per female as a measure of fecundity. Thus, we obtained female fecundity after 6 hours, 1 day and 4 days of exposure to flour conditioned by either MB or FB groups. In a second experiment, we conditioned the flour using either female-only or male-only groups of beetles (three 7-day-old virgin beetles of the same sex). We then introduced a male and female (7-day-old virgins) into 2g of this conditioned flour for either 6 hours or 24 hours (*n* = 11-12 pairs per conditioning treatment per duration of exposure). Following this, we estimated the number of eggs laid by each female as described above.

### Testing the impact of stink gland contents on female fecundity

Flour beetles have paired abdominal and thoracic stink glands that secrete a mix of chemicals including toxic quinones [26]. To test whether secreted products from beetles’ stink glands directly alter female fitness, we measured female fecundity after a brief exposure to abdominal stink glands. We paired 7-day-old virgins (1 male + 1 female) in 2 g flour for 2 days (*n* = 10 pairs per treatment). We separated the females and transferred them individually into a 35mm Petridish whose bottom was covered with a filter paper for traction, and a thin layer of flour to avoid starving the beetles during the assay. On the same day, we separately dissected abdominal stink glands from 60 MB and 60 FB females (sex ratio groups constituted as described earlier). For each set, we used a micro-pestle to pool and homogenize the glands in 300μl hexane, and centrifuged the suspension (5000 rpm, 5 minutes). 10μl of the supernatant was thus equivalent to the abdominal gland content of 2 females. We then preformed serial dilution of the supernatant to obtain abdominal gland content equivalents of 0.75, 0.5, 0.25, 0.1, 0.05, 0.025 and 0.001 females in a total volume of 10μl (*n* = 5-6/ dilution/sex ratio). We soaked filter paper discs (10 mm diameter) with 10μl of the supernatant (or hexane as control), kept them under laminar air flow for 10 minutes to evaporate the solvent, and then placed the discs at the center of a Petriplate containing the experimental female. Twelve hours later, we tested the fecundity of each female as described above.

Next, we wanted to identify the chemical component of female stink glands responsible for reduced female fecundity. Previous analyses show that ethylbenzoquinone (EBQ) and methylbenzoquinone (MBQ) are two major components of *T. castaneum* stink glands [27]. Hence, we quantified the amount of EBQ and MBQ produced by females in MB and FB groups. We constituted MB and FB groups as earlier (*n* = 10 groups per sex ratio), and after 7 days we dissected abdominal and thoracic stink glands of a randomly chosen female from each group. For each female, we combined abdominal and thoracic gland samples and homogenized in 60μl cold methanol. Subsequently, we quantified the amount of EBQ and MBQ in each sample using HPLC (see SI methods for details). For calibration, we used pure MBQ (Sigma) and lab-synthesized EBQ (see SI Methods).

Finally, to directly test whether quinones regulate fecundity, we exposed 9 day-old test females (reared as a single mating pair for 2 days before exposure, as described above) to abdominal stink gland extracts of MB females containing varying concentrations of added EBQ and MBQ. We added 10μl of the gland extract equivalent to 0.5 females, supplemented with 1, 5 or 10 μg of either quinone to a filter paper disc (10 mm diameter), and placed the disc in a 35 mm Petridish containing a female (*n* = 7-10 females per concentration per chemical). As controls, we exposed test females to discs spotted only with solvent, or with the MB gland extract without quinones. After exposing females to each chemical mixture for 12 hours, we measured fecundity as described earlier.

## RESULTS

### Male-biased sex ratio increases female fitness

We found that females from male biased (MB) groups outperformed females from unbiased (UB) and female biased (FB) groups. Throughout their major reproductive lifespan, MB females consistently produced more offspring compared to UB or FB females (Fig 1A; Table S1), resulting in over twice as many total offspring per female (Fig 1B; Table S1). The differential reproductive success was driven by offspring quantity: MB females laid ~three times more eggs per female per day, compared to UB or FB females (Fig 1C; Table S1). We observed a similar pattern even when we used a larger group size (FB: 9 females + 3 males, and MB: 3 females + 9 females; Fig S1). Thus, a male-biased sex ratio increased female reproductive fitness by increasing female fecundity. We also found significantly fewer deaths in MB females, compared to both UB and FB females (Fig. S2A; Table S1; Fisher’s exact test, two tailed p value: MB vs. FB = 0.009; MB vs. UB = 0.001; UB vs. FB = 0.999). In FB replicates where a female died earlier in the experiment, the sex ratio would become less skewed, leading us to underestimate the impact of the skewed sex ratio. In contrast, female mortality in UB groups would increase the skew in sex ratio. Despite these confounding factors, female fertility and fecundity in FB and UB groups was consistently lower than MB groups, suggesting that early effects of sex ratio were very strong. MB females also had better immune function (Fig S2B-D; Table S1). Together, these data indicate that low fecundity of FB and UB females cannot be explained by tradeoffs with other fitness components. In subsequent experiments, we focused on the mechanism underlying the observed impact of sex ratio on female reproductive fitness.

**Figure 1:**
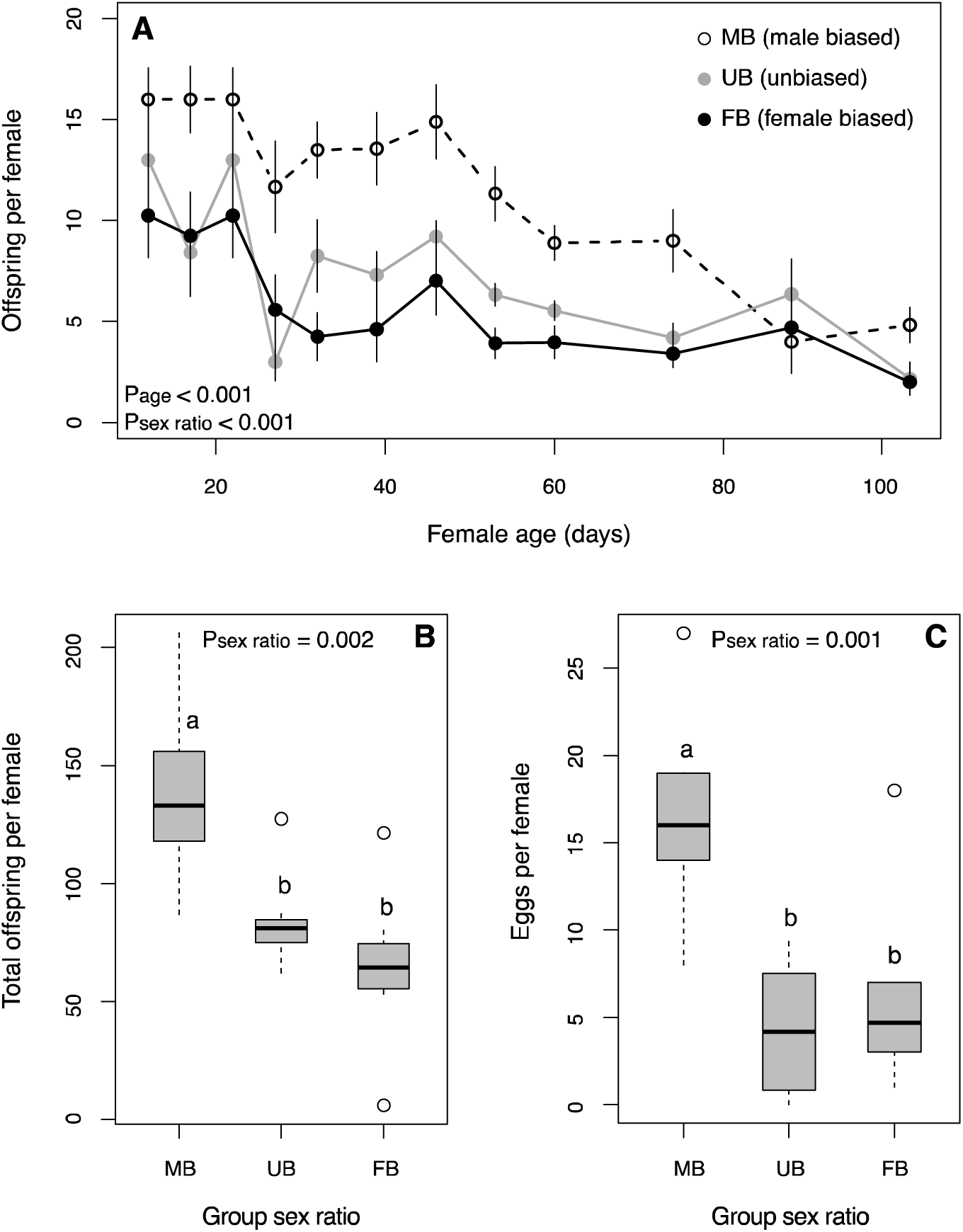
Female fitness as a function of group sex ratio. (A) Average offspring (±se) produced per female as a function of age (days post eclosion) (B) Total offspring produced per female during the experiment and (C) Fecundity of 28-day-old females as a function of sex ratio. MB = male-biased groups (3 males + 1 female); UB = unbiased groups (3 males + 3 females); FB = female-biased groups (1 male + 3 females). Box plots show median and quartiles, and lowercase alphabets indicate significantly different groups inferred from pairwise comparisons.

### Sexual interactions do not explain female fitness

We evaluated the hypothesis that female fitness increases when females can selectively choose to mate with superior males. This hypothesis predicts that FB females should have low fitness (no mate choice possible), and UB and MB females should have equally higher fitness (choice between 3 males in each case). Instead, we found that females in both FB and UB groups had lower fitness than females in MB groups (Fig 1). Thus, increased opportunity for female mate choice was unlikely to explain the higher fitness of MB females. An alternative hypothesis is that male mate choice drives female fitness. This hypothesis predicts that within replicates of UB and FB groups, only one female would have high fitness. However, when we only considered the female with the highest fitness in each group, we still found that MB females outperformed FB and UB females (Fig S2E). Thus, it is unlikely that opportunity for male or female mate choice was responsible for the increased fitness of MB females. Nonetheless, we specifically tested the impact of sex ratio on mating behaviour and female fecundity. We found that although MB females received more matings and tended to spend more time in copula (Fig S3, Table S2), female fitness was not correlated with either of these behaviours (Fig 2). In other words, adding mating behaviours to a model with group sex ratio as an explanatory variable did not improve the model fit (Table S2). Group sex ratio did not affect other aspects of mating behaviour (Fig S3). Thus, sexual interactions between individuals could not explain the higher fitness of MB females.

**Figure 2:**
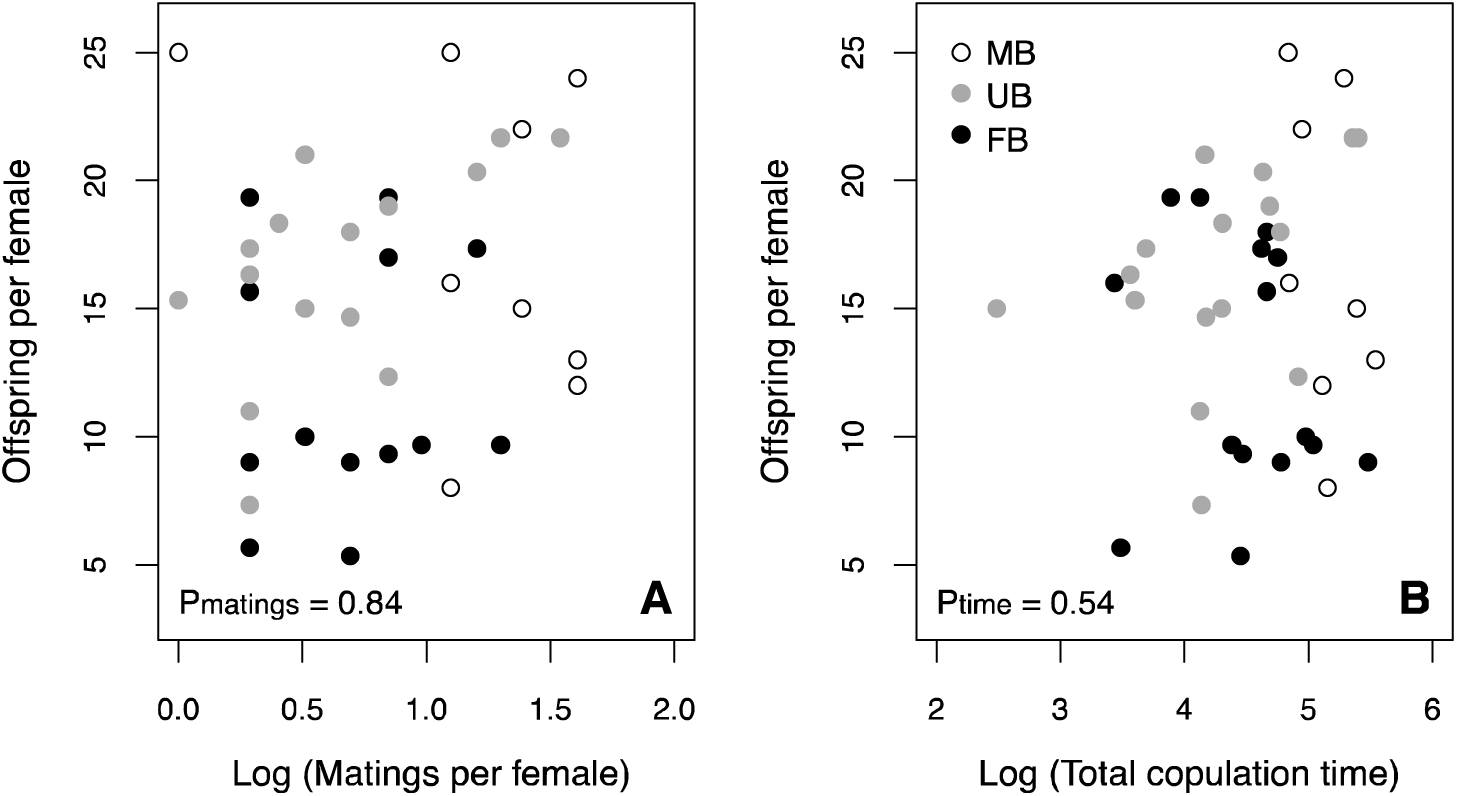
Female reproductive fitness as a function of mating behaviour. Average offspring per female, as a function of (A) Total number of matings per female in a 30-min observation period (B) Total time spent in copula per female. The x-axis was log-transformed in both panels. MB = male-biased groups (3 males + 1 female); UB = unbiased groups (3 males + 3 females); FB = female-biased groups (1 male + 3 females).

### Female-secreted benzoquinones explain the fitness impact of sex ratio

Flour beetles secrete toxic compounds in the flour, which are suggested to decrease female fecundity at high concentrations [28,29]. Thus, it is possible that greater accumulation of toxins in flour used by FB groups was responsible for the decreased fecundity of FB females. To test this, we measured the reproductive output of mated females exposed to flour previously “conditioned” by either FB or MB groups. We found that females responded rapidly to cues in conditioned flour, with a large reduction in fecundity after only 6 hours of exposure to flour conditioned by FB groups (Fig 3A, Fig S4A; Table S3). Critically, flour conditioned by females also produced a similar decline in fecundity (Fig 3B, Fig S4B; Table S3). These results indicated that the reduced fitness of FB females was mediated by female secreted chemicals in the flour, rather than via direct inter- or intra-sexual interactions.

**Figure 3:**
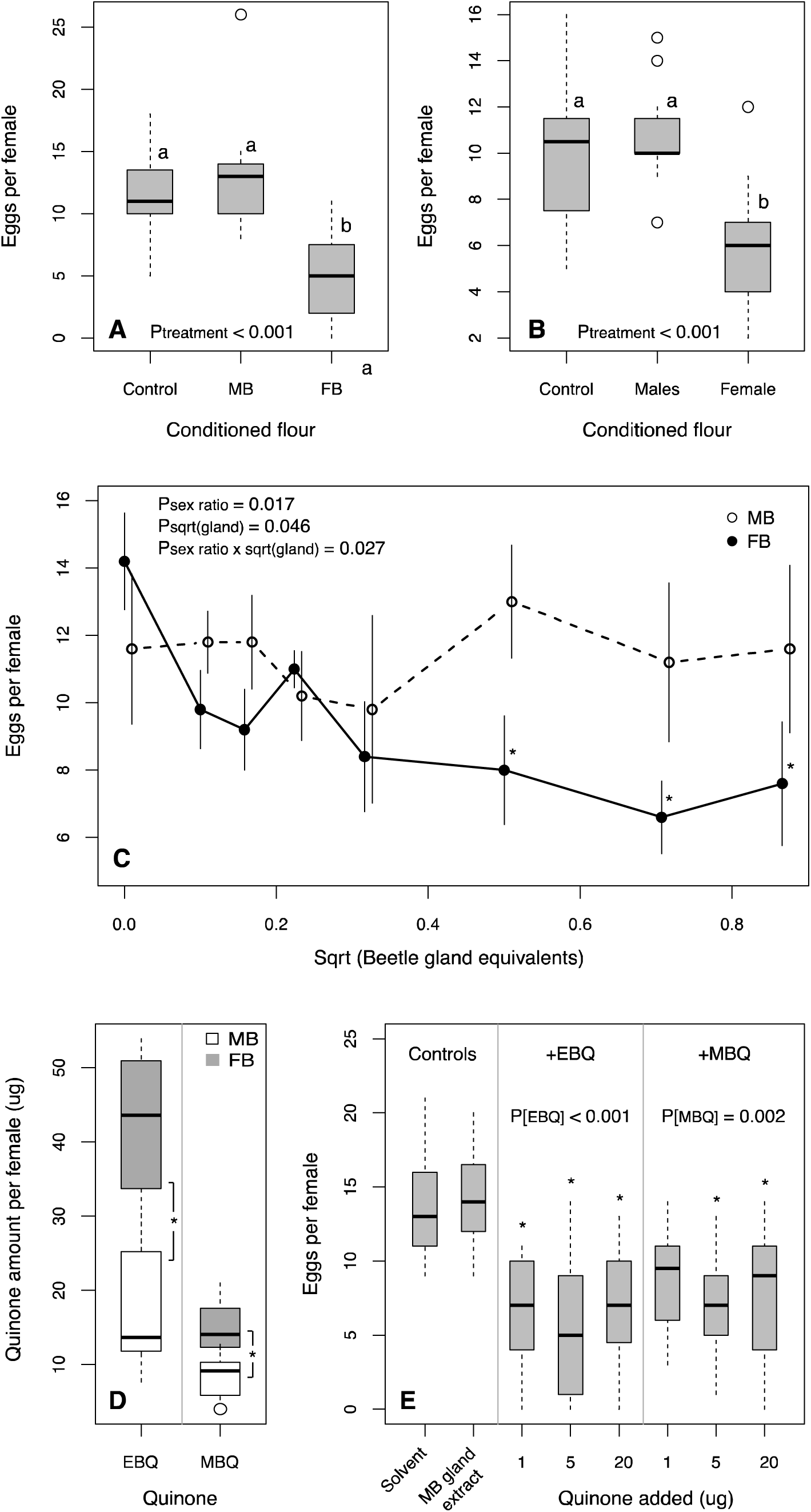
Quinones in female stink glands mediate fitness effects of group sex ratio. Fecundity of test females exposed to flour conditioned by (A) MB vs. FB groups, or (B) virgin females vs. males, for 6 hours (controls: unconditioned flour). Lowercase alphabets indicate significantly different groups inferred from pairwise comparisons. (C) Mean fecundity (±se) of test females exposed to increasing concentrations of stink gland extracts of females from MB vs. FB groups. MB data are slightly displaced along the x-axis for clarity. Asterisks indicate treatments that are significantly different from the respective control for each treatment (concentration 0). (D) Amount of ethyl- and methyl-benzoquinones (EBQ and MBQ) in stink glands of females from MB vs. FB groups. (E) Fecundity of test females exposed to MB stink gland extracts containing varying amounts of either EBQ or MBQ (controls: solvent used for gland extraction, and MB gland extract without added quinones). Box plots show median and quartiles; asterisks indicate groups that are significantly different from the control (MB gland extract).

Previous work shows that female flour beetles secrete more quinones (produced by stink glands) than males [25,27]. Hence, we tested whether stink gland secretions (rather than other secretions or excreta in the flour) were responsible for the observed effects of female-conditioned flour on female fitness. We found that fecundity of previously mated test females decreased after exposure to abdominal stink gland contents of FB females, whereas gland contents of MB females did not alter fecundity (Fig 3C; Table S3). These results confirmed that the contents of female stink glands directly affected female fitness. Next, we measured the concentration of benzoquinones (ethyl benzoquinone, EBQ and methyl benzoquinone, MBQ) in abdominal and thoracic stink glands dissected from females from MB vs. FB groups. We found that stink glands of FB females had higher amounts of both EBQ and MBQ (Fig 3D; Table S3), suggesting that these quinones may be responsible for lower fecundity of FB females. Finally, we tested whether adding quinones to gland extracts of MB females could mimic the effects of glands from FB females. Indeed, we found that adding EBQ or MBQ to stink gland extracts of MB females reduced the fecundity of test females (Fig 3E; Table S3).

Together, our results show that at high female density (i.e. in UB and FB groups), females upregulate EBQ and MBQ production, suppressing oviposition by all females in the group and resulting in low female fitness. In contrast, MB females secrete less MBQ and EBQ, allowing MB females to maximize oviposition and fitness. We infer that the observed female response to high female density reflects female interference competition for shared dietary and oviposition resources.

## DISCUSSION

A rich body of theoretical and empirical work documents the varied causes and consequences of adult sex ratio for individual fitness. Perhaps due to the obvious implications for sexual conflict and sexual selection, the large majority of this work has been interpreted in light of female mate choice and male male intrasexual competition. However, we found that neither male mate competition nor increased opportunity for female mate choice explained higher female fitness in male-biased groups of flour beetles. Instead, female competitive interactions were responsible for the observed fitness difference between male-biased and female-biased groups, regardless of group size. Similarly, in houseflies a male-biased sex ratio increased female fecundity and offspring survival [30]. Although male houseflies courted more frequently in male biased groups, female fitness was not associated with male courtship and was not affected by the opportunity for female mate choice. As discussed by the authors, these results suggest cryptic effects of social, non-sexual interactions between individuals as drivers of increased female fitness under a male biased sex ratio. Such non-sexual competitive interactions between females may be widespread, with multiple examples in mammals [reviewed in 13] as well as in social insects (discussed in [31]) and mosquitofish [32,33]. Thus, non-sexual interactions between females may frequently mediate the outcome of altered sex ratio in multiple taxa.

Importantly, we identified female-secreted benzoquinones as mediators of female competitive interactions in flour beetles. Studies on *Tribolium* have long implicated quinones as a mechanism of negative density-dependent population growth [34–36], though this was never explicitly tested. Quinones are secreted from abdominal and thoracic glands [26], and accumulate in flour in a density-dependent manner [16]. At high concentrations, quinones are toxic, causing developmental abnormalities in juveniles [16], and may thus represent a form of interference competition at high density. Indeed, beetles typically avoid flour containing high amounts of quinones [37]. The individual response to major quinone components also increases as a function of population density [38]. For instance, exposing a pair of focal beetles to olfactory cues (primarily quinones) from crowded beetles for three days decreased female oviposition rate by up to 25% [36]. Therefore, we hypothesized that quinones might be responsible for the observed differences in female fitness as a function of sex ratio. If true, the quinone hypothesis would predict that MB groups should produce less quinones than FB or UB groups, and exposure to more quinones should decrease fecundity. We found support for both these predictions in independent experiments: MB females had less quinones in their stink glands, and adding quinones to gland extracts of MB females mimicked the fitness impact of gland extracts of FB females. Our results also suggest that females are particularly sensitive to conditioned flour, and respond by modulating their fecundity. Previous experiments support this finding: under a balanced sex ratio, net female fecundity decreased rapidly with increasing population density, but the decline was slower in highly male biased groups containing a single female [39]. Previous work also shows that sensitivity to conditioned flour is a heritable trait that can rapidly evolve under selection in laboratory populations, and is controlled by a single (or few) genes [40]. Thus, density dependence itself may evolve via sensitivity to quinones. Higher quinone sensitivity in females may also explain our observation that male mortality was not affected by group sex ratio, even though males were exposed to high quinone levels in FB and UB groups.

Could quinones also account for the observed impact on other fitness proxies such as lifespan and immune traits? Quinones are toxic for *T. castaneum* juveniles [16], and cause high adult mortality in related species such as *T. destructor* [41]. A recent study showed that beetles grew faster and larger in male-biased groups [42], possibly due to greater amounts of toxic quinones in female-biased groups. Thus, it is fairly straightforward to explain the lower lifespans of UB and FB females. However, it is more difficult to explain the mechanism responsible for poor immune function in these females. One possibility is that similar to the impact on lifespan, quinone toxicity may decrease overall body condition, resulting in poor immune responses. Alternatively, quinone production may trade off with innate immunity. However, these possibilities need to be explicitly tested. Although our results suggest that quinones accumulate in the flour in proportion to their concentration in stink glands, further work is necessary to confirm this and to test whether females may also control the secretion rate of stink gland contents in a density-dependent manner. The precise mechanism through which benzoquinones suppress fecundity also remains to be elucidated; possibilities include direct effects on egg maturation or oviposition behaviour, or indirect effects via physiological costs of producing quinones.

Our finding that female fitness increases as a function of male biased sex ratio contradicts some previous work with *T. castaneum* showing inter-sexual conflict. Male competitive ability and mating success increased significantly in experimentally evolved male-biased populations [43]. Increasing the number of males in mating trials decreased the fitness of evolved FB females, but not of evolved MB females. As the authors discuss, these data suggest that females became more sensitive to (as yet unknown) deleterious effects of multiple mating, indicating underlying sexual conflict. However, unlike *Drosophila* where sexual conflict is mediated via mating-related harm to females (discussed in [43]), there is no direct evidence for such harm in *T. castaneum.* Furthermore, the impact of female-female interactions in the evolved FB lines is unclear. Testing quinone production and sensitivity in the evolved male-biased and female-biased lines may help to understand the conflicting results. In contrast, two other studies with *T. castaneum* corroborate our results, showing increasing female fitness with increasing male-biased sex ratios. First, populations selected for female-biased sex ratios showed a reduction in fitness [44]. Second, Wade measured the total number of offspring produced by groups with sex ratios ranging from 0.2 to 0.8 after 50 days [45]. Re-analyzing these data, we found that per capita female fitness decreased as a function of the proportion of females in the group (Fig S5A). Similarly, in another set of experiments (Wade, unpublished data), per capita female productivity declined as a function of the number of females in groups with different sex ratios maintained at a constant density (Fig S5B). Despite the many differences in experimental design, these results support our findings and suggest that female resource competition may be the most important determinant of female fitness in *T. castaneum*.

In closing, we suggest that the role of females in the evolution and consequences of group sex ratio has generally been underappreciated so far [13]. This is unfortunate because sex ratios provide a context where both sexual selection and resource competition may be altered. Our work demonstrates that fitness consequences of male biased sex ratios in flour beetles are driven by chemically mediated female non-sexual competitive interactions. Specifically, we show that females use benzoquinones as a major weapon in this warfare, although the mechanism responsible for the resulting suppression of fecundity remains unclear. We suggest that such effects of intrasexual female interference competition may not be uncommon, and may frequently confound the effects of sex ratio on female fitness.

## AUTHOR CONTRIBUTIONS

IK and DA conceived and designed experiments with input from other authors; IK, AP, SI, MU and PL conducted beetle experiments; RS, NJ and RV designed and conducted chemical synthesis and analyses; DA and IK analyzed data and wrote the manuscript with input from other authors.

## ACKNOWLEDGEMENTS

We thank Michael Wade for sharing his unpublished data, and Kavita Isvaran for insightful discussions. We acknowledge funding and support from the National Centre for Biological Sciences, SERB-DST Young Scientist Program (IK), University Grants Commission and Council for Scientific and Industrial Research (MU), and the DST INSPIRE Faculty Program (DA).

## REFERENCES

[1] Vlad MO. The optimal sex ratio for age-structured populations. Mathematical Biosciences 1989;93:181–90. doi:10.1016/0025-5564(89)90022-9.

[2] Emlen ST, Oring LW. Ecology, sexual selection, and the evolution of mating systems. Science 1977;197:215–23.

[3] Fisher RA. The Genetical theory of natural selection. The Clarendon Press; 1930.

[4] Kvarnemo C, Ahnesjo I. The dynamics of operational sex ratios and competition for mates. Trends in Ecology & Evolution 1996;11:404–8.

[5] Nandy B, Gupta V, Sen S, Udaykumar N, Samant MA, Ali SZ, et al. Evolution of mate-harm, longevity and behaviour in male fruit flies subjected to different levels of interlocus conflict. BMC Evol Biol 2013;13:1–1. doi:10.1186/1471-2148-13-212.

[6] Holland B, Rice WR. Experimental removal of sexual selection reverses intersexual antagonistic coevolution and removes a reproductive load. Proc Natl Acad Sci USa 1999;96:5083–8.

[7] Sakurai G, Kasuya E. The costs of harassment in the adzuki bean beetle. Animal Behaviour 2008;75:1367–73. doi:10.1016/j.anbehav.2007.09.010.

[8] Stockley P. Sexual conflict resulting from adaptations to sperm competition. Trends in Ecology & Evolution 1997;12:154–9.

[9] Berglund A. The operational sex ratio influences choosiness in a pipefish. Behavioral Ecology 1994;5:254–8. doi:10.1093/beheco/5.3.254.

[10] Arnqvist G, Nilsson T. The evolution of polyandry: multiple mating and female fitness in insects. Animal Behaviour 2000;60:145–64. doi:10.1006/anbe.2000.1446.

[11] West-Eberhard MJ. Sexual selection, social competition, and speciation. Q Rev Biol 1983;58:155–83. doi:10.2307/2828804?ref=no-x-route:4fb252f28a89822717b8cfaa73c307f8.

[12] Tobias JA, Montgomerie R, Lyon BE. The evolution of female ornaments and weaponry: social selection, sexual selection and ecological competition. Philos Trans R Soc Lond, B, Biol Sci 2012;367:2274–93. doi:10.1098/rstb.2011.0280.

[13] Stockley P, Bro-Jørgensen J. Female competition and its evolutionary consequences in mammals. Biological Reviews 2011;86:341–66. doi:10.1111/j.1469-185X.2010.00149.x.

[14] Bell MBV, Cant MA, Borgeaud C, Thavarajah N, Samson J, Clutton-Brock TH. Suppressing subordinate reproduction provides benefits to dominants in cooperative societies of meerkats. Nat Commun 2014;5:4499. doi:10.1038/ncomms5499.

[15] Harvey JA, Poelman EH, Tanaka T. Intrinsic inter- and intraspecific competition in parasitoid wasps. Annu Rev Entomol 2013;58:333–51. doi:10.1146/annurev-ento-120811-153622.

[16] Sokoloff A. The Biology of *Tribolium*. London: Oxford University Press; 1977.

[17] Lewis SM, Kobel A, Fedina T, Beeman RW. Sperm stratification and paternity success in red flour beetles. Physiol Entomol 2005;30:303–7. doi:10.1111/j.1365-3032.2005.00450.x.

[18] Fedina TY, Lewis SM. Female mate choice across mating stages and between sequential mates in flour beetles. J Evolution Biol 2007;20:2138–43. doi:10.1111/j.1420-9101.2007.01432.x.

[19] Fedina TY. Cryptic female choice during spermatophore transfer in *Tribolium castaneum* (Coleoptera: Tenebrionidae). J Insect Physiol 2007;53:93–8. doi:10.1016/j.jinsphys.2006.10.011.

[20] Arnaud L, Haubruge E. Mating behaviour and male mate choice in *Tribolium castaneum* (Coleoptera, Tenebrionidae). Behaviour 1999;136:67–77. doi:10.1163/156853999500677.

[21] Pai A, Feil S, Yan G. Variation in polyandry and its fitness consequences among populations of the red flour beetle, *Tribolium castaneum*. Evol Ecol 2007;21:687–702. doi:10.1007/s10682-006-9146-4.

[22] Lewis SM, Austad SN. Sources of intraspecific variation in sperm precedence in red flour beetles. Am Nat 1990;135:351–9. doi:10.2307/2462251?ref=no-x-route:0ec1a02474599c03350950cfc611e982.

[23] Bernasconi G, Keller L. Female polyandry affects their sons’ reproductive success in the red flour beetle *Tribolium castaneum*. J Evolution Biol 2001;14:186–93. doi:10.1046/j.1420-9101.2001.00247.x.

[24] Parent CE, Agashe D, Bolnick DI. Intraspecific competition reduces niche width in experimental populations. Ecol Evol 2014;4:3978–90. doi:10.1002/ece3.1254.

[25] Khan I, Prakash A, Agashe D. Immunosenescence and the ability to survive bacterial infection in the red flour beetle *Tribolium castaneum*. Journal of Animal Ecology n.d.

[26] Markarian H, Florentine GJ, Pratt JJ Jr. Quinone production of some species of *Tribolium*. J Insect Physiol 1978;24:785–90. doi:10.1016/0022-1910(78)90096-3.

[27] Unruh LM, Xu R, Kramer KJ. Benzoquinone levels as a function of age and gender of the red flour beetle, *Tribolium castaneum*. Insect Biochemistry and Molecular Biology 1998;28:969–77. doi:10.1016/S0965-1748(98)00085-X.

[28] Park T. Studies in population physiology. III. The effect of conditioned flour upon the productivity and population decline of *Tribolium confusum*. J Exp Zool 1934;68:167–82. doi:10.1002/jez.1400680202.

[29] Park T. Studies in population physiology. VI. The effect of differentially conditioned flour upon the fecundity and fertility of Tribolium confusum Duval. Journal of Experimental Zoology 1936;73:393–404. doi:10.1002/jez.1400730303.

[30] Carrillo J, Danielson-François A, Siemann E, Meffert L. Male-biased sex ratio increases female egg laying and fitness in the housefly, *Musca domestica*. J Ethol 2011;30:247–54. doi:10.1007/s10164-011-0317-6.

[31] Berglund A, Magnhagen C, Bisazza A, König B, Huntingford F. Female-female competition over reproduction. Behavioral Ecology 1993;4:184–7.

[32] Smith CC, Sargent RC. Female fitness declines with increasing female density but not male harassment in the western mosquitofish, *Gambusia affinis*. Animal Behaviour 2006;71:401–7. doi:10.1016/j.anbehav.2005.06.003.

[33] Smith CC. Independent effects of male and female density on sexual harassment, female fitness, and male competition for mates in the western mosquitofish *Gambusia affinis*. Behav Ecol Sociobiol 2007;61:1349–58. doi:10.1007/s00265-007-0365-7.

[34] Park T. Experimental studies of insect populations. Am Nat 1937;71:21–33. doi:10.2307/2457241?ref=no-x-route:9d7c3751097eb3a2b02021eef95d46cd.

[35] Park T, Woollcott N. Studies in population physiology. VII. The relation of environmental conditioning to the decline of *Tribolium confusum* populations. Physiological Zoology 1937; 10:197–211. doi:10.2307/30160900?ref=no-x-route:191b77aa5b7796aeddbf693a50343bca.

[36] Sonleitner FJ, Gutherie PJ. Factors affecting oviposition rate in the flour beetle *Tribolium castaneum* and the origin of the population regulation mechanism. Researches on Population Ecology 1991;33:1–11. doi:10.1007/bf02514569.

[37] Loconti JD, Roth LM. Composition of the odorous secretion of *Tribolium castaneum*. Annals of the Entomological Society of America 1953;46:281–9. doi:10.1093/aesa/46.2.281.

[38] Duehl AJ, Arbogast RT, Teal PEA. Density-related volatile emissions and responses in the red flour beetle, *Tribolium castaneum*. J Chem Ecol 2011;37:525–32. doi:10.1007/s10886-011-9942-3.

[39] Birch LC, Park T, Frank MB. The effect of intraspecies and interspecies competition on the fecundity of two species of flour beetles. Evolution 1951;5:116–32. doi:10.2307/2405763?ref=no-x-route:712b75169862515404fdcd8d9a518ce4.

[40] Lavie B, Ritte U, Moav R. The genetic basis of egg lay response to conditioned medium in the flour beetle, *Tribolium castaneum*: I. Two-way selection. Theor Appl Genet 1978;52:193–9. doi:10.1007/BF00273889.

[41] Palm N-B. Structure and physiology of the stink glands in *Tribolium destructor* Uytt. Opusc Ent 1946;11:119–32.

[42] Ellen ED, Peeters K, Verhoeven M, Gols R, Harvey JA, Wade MJ, et al. Direct and indirect genetic effects in life-history traits of flour beetles *(Tribolium castaneum)*. Evolution 2016;70:207–17. doi:10.1111/evo.12835.

[43] Michalczyk Ł, Millard AL, Martin OY, Lumley AJ, Emerson BC, Gage MJG. Experimental evolution exposes female and male responses to sexual selection and conflict in *Tribolium castaneum*. Evolution 2010;65:713–24. doi:10.1111/j.1558-5646.2010.01174.x.

[44] Lavie B, Beiles A. Different types of response to selection for sex ratio in the flour beetle *Tribolium castaneum*. Ann Genet Sel Anim 1981;13:119–30. doi:10.1186/1297-9686-13-2-119.

[45] Wade MJ. Variance-effective population number: The effects of sex ratio and density on the mean and variance of offspring numbers in the flour beetle, *Tribolium castaneum*. Genet Res 1984;43:249–56.

